# Intracellular FUS protein accumulation leads to cytoskeletal, organelle and cellular homeostasis perturbations

**DOI:** 10.1101/2022.10.04.510756

**Authors:** Chyi Wei Chung, Alexandra J. Zhou, Ioanna Mela, Amberley D. Stephens, Akinori Miyashita, Peter H. St George-Hyslop, Clemens F. Kaminski, Tuomas P. J. Knowles, Gabriele S. Kaminski Schierle

## Abstract

The molecular mechanisms that connect the formation of aberrant cytoplasmic FUS condensates to biological malfunction are incompletely understood. Here, we develop an approach to determine the intracellular FUS viscosity in live mammalian cells and find that ALS-related mutant P525L-FUS forms the most viscous condensates and has impaired cytoskeletal mechanoproperties and increased euchromatin formation. We further show that some of the main cellular organelles, e.g., actin/tubulin, lysosomes, mitochondria, the endoplasmic reticulum, are significantly functionally/structurally impaired in the presence of FUS. These may be related to defects in the tubulin network, as the latter facilitates transport, formation, fusion and fission of organelles. We observe significant increases in lysosomal biogenesis, size and pH; moreover, intracellular FUS accumulation significantly promotes cytoplasmic-to-nuclear translocation of TFEB, i.e., the master gene for inducing autophagy. However, despite these, increased autophagy needed for protein aggregate clearance is not observed to occur. Our study reveals that the formation of highly viscous FUS condensates significantly impacts cytoskeletal/organelle function and cellular homeostasis, which are closely associated with cell ageing. This raises the intriguing question as to whether mutant FUS activates similar cell processes as those during cellular senescence.

## Introduction

The aberrant condensation of fused-in sarcoma (FUS) is a hallmark of both amyotrophic lateral sclerosis (ALS) and frontotemporal dementia (FTD). Under physiological conditions, FUS undergoes liquid-liquid phase separation (LLPS) and exists in a liquid and/or a dense condensate state predominantly within cell nuclei. Prominent cytoplasmic mislocalisation and the formation of more viscous, and in many cases, gelled condensates or even fibrillar aggregates, are observed upon missense and truncation mutations. Many, but not all of these ALS-associated mutations occur within the FUS nuclear localisation signal (NLS) (Bosco et al., 2010; Conte et al., 2012; Waibel et al., 2010). Pathological phase separation and the formation of gelled condensates have also been linked to arginine hypomethylation of FUS, as commonly observed in cases of sporadic FTD, which leads to disruptions in cytoplasmic ribonucleoprotein (RNP) granule function (e.g., stress granules (SGs) and neuronal transport granules) (Hofweber et al., 2018; Murakami et al., 2015; Qamar et al., 2018).

Whilst much attention has been given to the pathological role of FUS in the formation of cytoplasmic SGs, an understanding of how FUS condensation leads to disturbances in cellular homeostasis and function would be invaluable. This knowledge could also yield avenues for therapeutic intervention in diseases associated with the aberrant assembly of these proteins, such as FUS-associated ALS (fALS-FUS) and frontotemporal lobar degeneration (FTLD-FUS).

A potentially powerful clue to the latter may come from a detailed analysis of different fALS-FUS and FTLD-FUS models. We thus first established different FUS expressing cell lines and stress models in HEK293T cells before comparing the changes in intracellular FUS viscosity of the various FUS models. Combining single particle tracking (SPT-) and fluorescence lifetime imaging microscopy (FLIM), we develop a method to quantify intracellular FUS viscosity without the need for an external sensor/tracer, and that is higher-throughput than conventional fluorescence recovery after photobleaching (FRAP). To understand how FUS condensation leads to disturbances in cellular homeostasis, we then focused our analysis on the role of FUS in the nucleus and the cytoplasm, including its effect on the cytoskeleton and on organelle positioning and function. We show that P525L-FUS cells form the most viscous condensates, which affect the level of euchromatin formation, cytoskeletal proteins and organelle positioning and function, including lysosomes, mitochondria and the endoplasmic reticulum (ER). We show that the latter is linked to the observed lysosomal de-acidification, which acts as a trigger for the nuclear to cytoplasmic translocation of transcription factor EB (TFEB), i.e., the master modulator of autophagic genes. Instead of the expected upregulation of macroautophagy (henceforth referred to as autophagy), however, autophagy is blocked at an early stage, which promotes cellular malfunction and disease progression (Aman et al., 2021; Martini-Stoica et al., 2016).

## Results

### Fluorescence lifetime imaging and single particle tracking form a tool to measure FUS condensate viscosity in live cells

Studying phase separation in live cells, and thus characterising intracellular FUS viscosity is challenging (Kuimova, 2012). We have previously shown, however, that protein phase separation and aggregation can be measured *in vitro* and in live cells using FLIM (Chen et al., 2017; Esbjörner et al., 2014). We thus first investigated whether SPT, which permits exact viscosity measurements *in vitro*, can be correlated with FLIM. We generated LLPS condensates by mixing equal amounts of recombinant GFP- and unlabelled WT-FUS protein. We then added the protein mixture into a solution containing 10% polyethylene glycol 35 (PEG-35) and 150 mM potassium chloride (KCl) to mimic the molecular crowding and ionic environments of the nuclear and cytoplasmic intracellular compartments. As we were interested in aggregation kinetics and changes in the micro-rheology of FUS with time, we performed ageing experiments involving SPT and FLIM over time, giving us a means to quantify both parameters of interest (**Figure 1A**). To probe the viscosity within FUS condensates, we introduced 40 nm fluorescent nanoparticles into the condensate mixture and tracked their trajectories. Because fluctuations seen in tracked nanoparticle trajectories are predominantly due to Brownian motion (Crocker and Grier, 1996); hence, we were able to calculate intra-condensate FUS viscosity from their mean square displacement (MSD) profiles. Between the 0 to 1h time points, FUS in the condensates becomes more viscous, with viscosity values increasing from 0.11±0.01 Pas to 0.44±0.08 Pas (**Figure 1B**, blue points). We observe a correlation between FLIM and SPT measurements (**Figure 1B**). During the same time period during which increased viscosity is observed, the GFP-tagged FUS molecules in the condensates cluster more closely, as indicated from the decrease in their fluorescence lifetimes from 2.23±0.01 ns to 2.08±0.01 ns (**Figure 1B**, red points; **Figure 1C**, Condensates). These values are in turn lower than that of dispersed WT-FUS before LLPS at 2.53 ns (**Figure 1B**, green point; **Figure 1C**, dispersed), which has a viscosity of 0.004 Pas from SPT measurements. To induce the fibrillation of FUS, we removed PEG-35 from the condensate mixture described above to yield solid fibrils with a fluorescence lifetime in the range of 2.01±0.01 ns (**Figure 1B**, magenta point; **Figure 1C**, Fibrils). As the fluorescence lifetime of GFP is independent of the viscosity of its surrounding environment (Davidson et al., 2020), we note that FLIM and SPT yield independent and complementary parameters indicative of the level of FUS compaction and viscosity, respectively. Our data further show that beyond an hour, there are no further significant changes in either fluorescence lifetime or viscosity.

**Figure 1:**
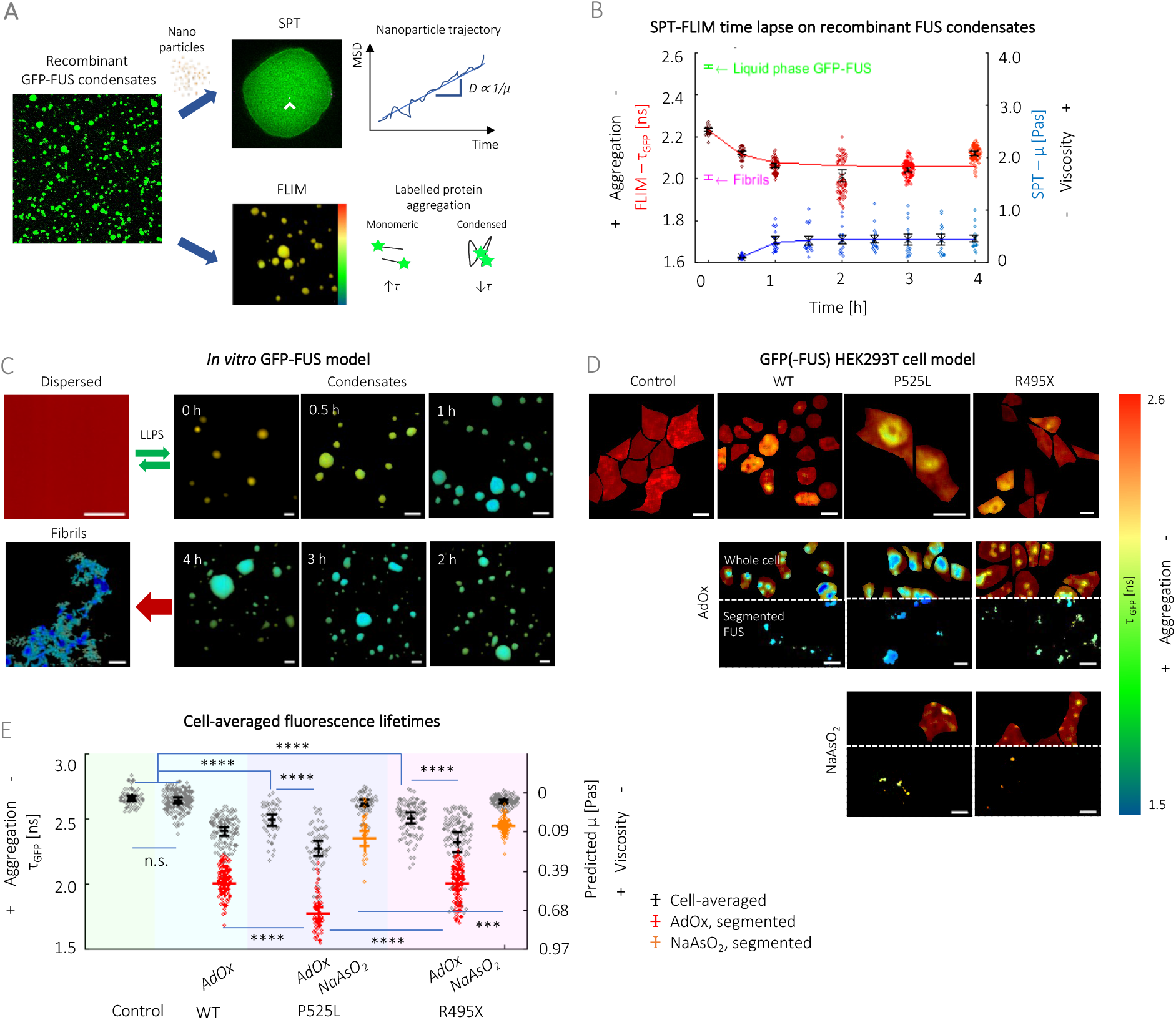
Fluorescence lifetime to viscosity calibration reveals that ALS-associated P525L-FUS forms the most viscous condensates. (A) FLIM and SPT measurements were performed on *in vitro* FUS condensates, giving a measure of condensation state and viscosity, respectively. For SPT, 40 nm nanoparticles (denoted by white arrowhead) were mixed into the condensates and the mean square displacement (MSD) profiles of their trajectory were analysed. (B) Combined SPT-FLIM data show that there is a trend between GFP fluorescence lifetimes (τ_GFP,_ FLIM in red) and viscosity (μ, SPT in blue), where more aged FUS condensates are also more viscous. Values plateau ∼1 hour and show no further change upon subsequent ageing. (C) Fluorescence lifetime maps for liquid-to-solid transition of recombinant FUS. A decrease in τ_GFP_ is used to quantify increasing condensation as dispersed FUS undergoes LLPS to form condensates, which are aged over a 4-hour period. Upon irreversible solid transition (i.e., formation of fibrils), fluorescence lifetime values are at the lowest values at 2.01±0.01 ns. Scale bars, 10 μm. (D) Fluorescence lifetime maps for FUS in the GFP HEK293T model, which show that fluorescence lifetimes fall in the same 1.5—2.6 ns range as the *in vitro* system in (C). The formation of hypomethylated (HYPO) intranuclear aggregates (first row, AdOx) and SGs (second row, NaAsO_2_) were induced by the addition of different stressors. These macro-assemblies can be easily segmented by fluorescence intensity thresholding (Segmented FUS). Scale bars, 10 μm. (E) Cell averaged-fluorescence lifetimes values (black) alongside viscosity predicted from correlative FLIM-SPT calibration on in vitro condensates (B). ALS-related FUS mutant cells have lower τ_GFP_ (i.e., are at a more condensed state) in comparison to WT-FUS and control cells. Fluorescence lifetimes of segmented hypomethylated nuclear aggregates (AdOx, red) and SGs (NaAsO_2_, orange) are also given, with P525L-forming more viscous macroassemblies compared to R495X-FUS. *In vitro* and HEK293T FUS cell measurements were based on 3 individual protein preparations and 3 biological repeats respectively. One-way ANOVA test (with Holm-Sidak’s multiple comparison), where n.s. is not significant, *** p<0.001 and **** p<0.0001.

To observe both physiological and aberrant condensate formation of FUS in a cell model, we created HEK293T cells expressing GFP-labelled FUS (**Figure S1A**). To ensure that our observations were not due to the accumulation of GFP, we included cells expressing a GFP only construct (i.e., Control) in our experiments. We studied WT-FUS, and two ALS-related mutants, (i.e., P525L and R495X with a point site mutation at and truncation of the NLS, respectively), thereby leading to cytoplasmic mislocalisation of FUS(Gonzalez et al., 2021). Pathological phase separation of FUS is also promoted upon hypomethylation of its arginine-rich domain, yielding irreversible FUS hydrogels that are prevalent in FUS-related forms of sporadic FTD (Hofweber et al., 2018; Murakami et al., 2015). In order to induce FUS hypomethylation, we treated the cells with adenosine periodate oxidase (AdOx), an arginine methyltransferase inhibitor, which modulates the methylation state of the RNA-binding, arginine-rich domain of FUS (Fujii et al., 2016; Qamar et al., 2018). This promotes gelation and results in the formation of clusters of hypomethylated (HYPO) intranuclear aggregates (**Figure S1B**). P525L-FUS cells lose cytoplasmic mislocalisation, whereas R495X-FUS cells (which have a truncated NLS) still retain cytoplasmic condensates after AdOx treatment. Furthermore, we induced the formation of SGs using sodium arsenite (NaAsO_2_) (**Figure S1C**). Existing as membraneless LLPS structures of cytoplasmic RNPs, SGs are transient structures of translationally stalled mRNA complexes and proteins (Sama et al., 2013). However, it has been suggested that in neurodegeneration models, they become persistent and are the site of aberrant aggregation (Wolozin and Ivanov, 2019). We validated the formation of SGs in our cell models by immunostaining of Ras-GTPase-activating protein binding protein (G3BP), a protein marker for the assembly and dynamics of SGs (**Figure S2**) (Yang et al., 2020). Agreeing with previous reports (Baron et al., 2013; Bosco et al., 2010), we observe colocalisation of FUS and G3BP only in the case of mutant variants of FUS (**Figure S2B**), despite the formation of SGs in all cell models.

### ALS-associated FUS mutants form the most viscous aggregates in live cells

Using FLIM, we could quantify the aggregation propensity and extent of FUS in our cell models (**Figure 1D&E**). We observe that the NLS-mutants have more aggregated FUS (cell-averaged values of 2.48±0.12 ns and 2.49±0.12 ns for P525L- and R495X-FUS, respectively), in comparison to nuclear-localised WT-FUS (2.62±0.07 ns) with fluorescence lifetimes that do not differ significantly from control cells at 2.64±0.06 ns. As FUS in SGs and hypomethylated intranuclear aggregates have higher fluorescence intensities than other compartments of the cell (**Figure S1B&C**), we could segment them by setting a fluorescence intensity threshold (**Figure 1D**, AdOx & NaAsO_2_). In comparison to R495X-, P525L-FUS has a lower fluorescence lifetime than hypomethylated intranuclear aggregates (1.78±0.05 ns *cf*. 2.01±0.04 ns in R495X) and SGs (2.35±0.06 ns *cf*. 2.46±0.02 ns in R495X). To calculate the intracellular viscosity of FUS, we created a fluorescence lifetime to viscosity calibration based on the *in vitro* FUS condensate system (**Figure S3A**), noting that GFP fluorescence lifetimes of recombinant FUS and the HEK293T FUS models fall within the same range of 1.6—2.6 ns (**Figure 1E**). Hence, we perform linear fitting (**Figure S3A**, black line) on SPT-FLIM data of the *in vitro* protein condensate system (**Figure S3A**, grey circles), to map fluorescence lifetime values of FUS imaged in live HEK293T cells to their corresponding viscosity values (**Figure 1E**, secondary axis). We note that before both AdOx and NaAsO_2_ treatment, the two NLS mutants of FUS are more viscous at ∼0.10 Pas than WT-FUS at 0.05 Pas. Moreover, we estimate that hypomethylated aggregates differ in their viscosity to SGs by a factor of 5—10, i.e., 0.69±0.05 (AdOx – P525L) to 0.14±0.06 Pas (NaAsO_2_ – P525L) and 0.47±0.04 (AdOx – R495X) to 0.04±0.02 Pas (NaAsO_2_ – R495X.

We observe a similar trend upon calculating the diffusion coefficient of these FUS macro-assemblies inside cells (**Figure S3B**). For the latter, we extracted MSD profiles of the FUS macro-assemblies’ trajectories over time, which we imaged in a similar manner to SPT measurements for the *in vitro* condensates. For both, AdOx and NaAsO_2_ treatments, P525L-FUS cells yield HYPO nuclear aggregates and SGs of lowest mobility, with respective diffusion coefficients of 0.036±0.004 nm^2^ s^-1^ (compared to 0.044±0.004 nm^2^ s^-1^ in WT-FUS cells) and 0.011±0.002 nm^2^ s^-1^ (compared to 0.025±0.001 nm^2^ s^-1^ in R495X-FUS cells). As aforementioned, R495X-FUS is the only model that retains the cytoplasmic distribution of FUS upon AdOx treatment due to its truncated NLS (**Figure S1B**). We observe that the hypomethylated aggregates it forms are the most mobile with a diffusion coefficient 0.076±0.007 nm^2^ s^-1^ (i.e., two-fold that of P525L- and WT-FUS). Hence, using both, our fluorescence lifetime to viscosity calibration as well as diffusion coefficient measurements, we show that there are differences in the way different FUS variants interact with their nuclear and cytoplasmic environment, which affect their aggregation propensity.

### P525L-FUS mutant impairs cellular mechanoproperties and enhances euchromatin formation

We have thus far established that there are differences in the condensation states of FUS, with mutant P525L- and R495X-FUS being in a more condensed state than WT-FUS, even if the latter has been exposed to stressors such as AdOx and NaAsO_2_. It has been suggested that loss of FUS functionality and its cytoplasmic mislocalisation can cause loss or gain of function defects by affecting essential cytoskeletal proteins (Theunissen et al., 2021). To further characterise the latter in our HEK293T cell models, we visualise filamentous (f-)actin and microtubules (i.e., the 2 main proteins of the cytoskeleton) using far-red fluorogenic Silicon Rhodamine (SiR) dyes (Lukinavičius et al., 2014) (**Figure 2**). As the SiR-dyes are of high specificity and only become fluorescent upon binding, their fluorescence intensity can be used as to measure f-actin and microtubule polymerisation levels. Our data reveal that only P525L- and not R495X-FUS has a significant impact on the cytoskeleton, as shown by reduction of f-actin, as shown by a loss in SiR-Actin intensity. A similar effect on SiR-Actin intensity can only be induced by stressors, such as AdOx and NaAsO_2_, which cause hypomethylation or SG formation, respectively (**Figure 2A&B**). However, in the case of P525L-FUS, the above stressors do not cause a further defect on the cytoskeleton as P525L-FUS alone does. We next used SiR-Tubulin and performed similar experiments as described above to see whether FUS condensates would also affect the microtubule protein, tubulin. It is observed that P525L-FUS, followed by R495X-FUS, cells have the lowest SiR-Tubulin intensity levels prior to drug treatments. Interestingly though, AdOx treatment, which promotes nuclear translocation of FUS and the formation of hypomethylated clusters, reduces the burden on microtubules mainly present in the cytoplasmic region, the effect being strongest in P525L-FUS cells, where the largest drop in SiR-Tubulin intensity has occurred prior to the addition of AdOx (**Figure 2C&D**). The opposite effect is seen when cytoplasmic SGs are formed using NaAsO_2_, which results in significantly reduced SiR-Tubulin intensities in all models. Additionally, we quantified the amounts of f-actin and β-tubulin (a sub-unit of microtubules) in the cell samples based on a Western Blot (**Figure S4**), and we find they do not differ significantly between FUS variants. Hence, this may indicate that depolymerisation of actin and microtubules has occurred, instead of lower concentrations of either being expressed.

**Figure 2:**
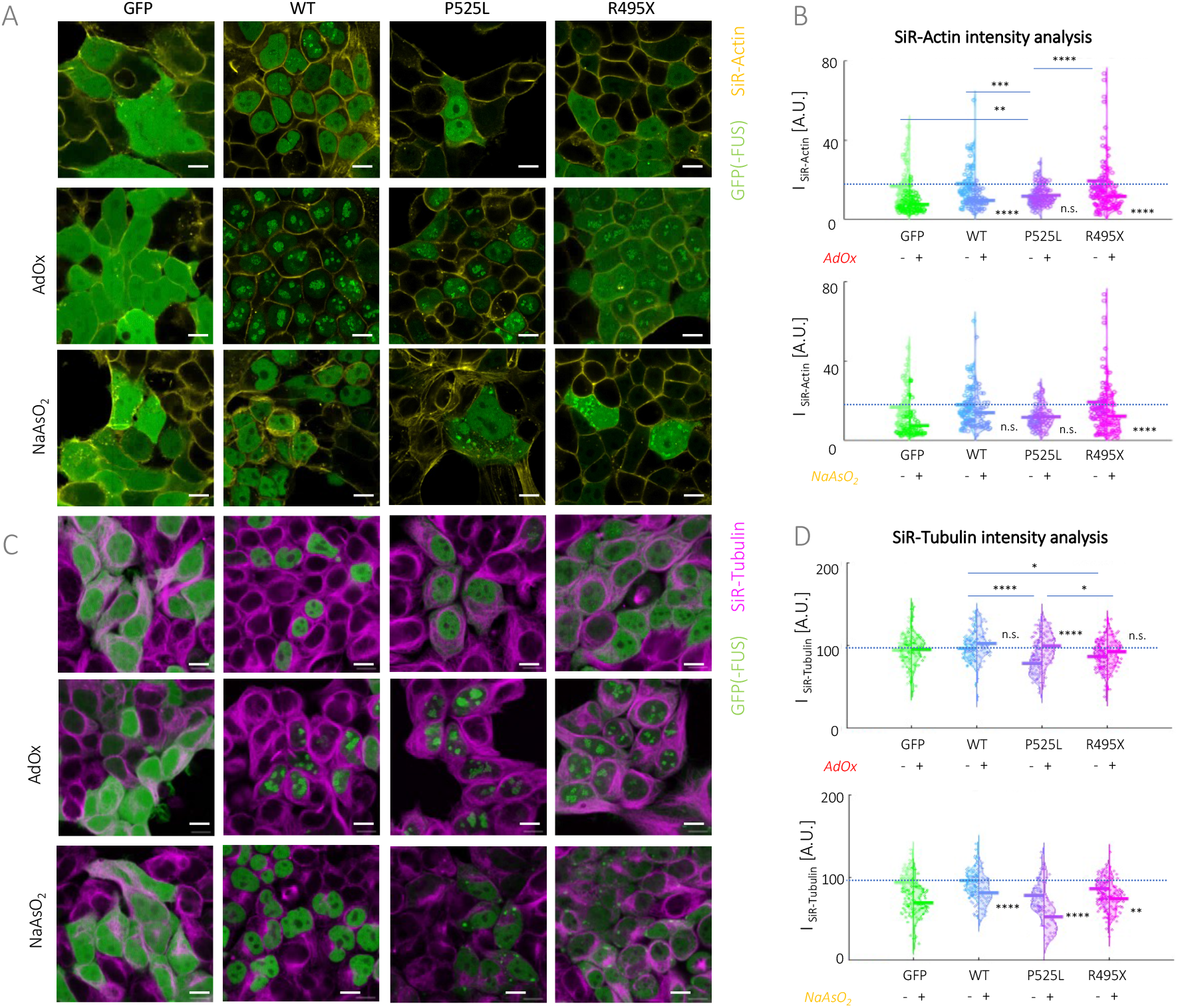
ALS-associated P525L-FUS has lowered levels of important cytoskeletal proteins based on SiR-dye staining. (A) Fluorescence intensity composite images for GFP(-FUS) (green) and SiR-Actin (yellow). Scale bars, 10 μm. (B) Cell-averaged fluorescence intensity quantification for SiR-Actin (I_SiR-Actin_). f-actin levels in P525L-FUS cells are low even without the addition of any stressors. AdOx has a greater effect in comparison to NaAsO_2_ on reducing f-actin levels. (C) Fluorescence intensity composite images for GFP(-FUS) (green) and SiR-Tubulin (magenta). Scale bars, 10 μm. (D) Cell-averaged fluorescence intensity quantification for SiR-Tubulin (I_SiR-Tubulin_). Tubulin levels in P525L- and R495X-FUS cells are impacted even without the addition of any stressors. The formation of HYPO aggregates improves microtubule levels, in contrast to SGs which impair microtubules. HEK293T measurements were based on 3 biological repeats. One-way ANOVA test (with Holm-Sidak’s multiple comparison), where * for p<0.05, ** p<0.01, *** p<0.001, **** p<0.0001.

To confirm that FUS condensates indeed affect cellular mechanoproperties by interfering with the two most important cytoskeletal proteins, actin and tubulin, we perform force-displacement (FD)-atomic force microscopy (AFM) measurements on the cytoplasmic regions of live cells (**Figure S5**). In comparison to WT-FUS cells, we have seen that mutant P525L-FUS cells display the lowest levels of polymerisation for both cytoskeletal proteins investigated; moreover, R495X-FUS cells have lower levels of SiR-Tubulin detected (**Figure 2**). Corroborating this, we find that P525L-FUS cells have the softest cytoplasm with an average apparent Young’s modulus (E_App_) of 12.3±2.2 kPa, followed by R495X at 13.7±4.0 kPa, in comparison to WT-FUS at 25.5±2.2 kPa (**Figure S5A**); these measurements correspond to intensity quantifications of SiR-dyes (**Figure 2**). Hence, we believe that the softening we observe is due to destabilisation of the cytoskeleton i.e., a decrease in E_App_ corresponds to lower intensities of the SiR-dyes (**Figure S5B**). It is interesting to note that the expression of a cytoplasmic protein, such as GFP in the control cells, also has an effect on the cell mechanoproperties as measured by FD-AFM. However, in contrast to FUS cells, which mainly leads to accumulation in the nucleus for WT-FUS and some cytoplasmic condensates in P525L- and R495X-FUS cells, no effect on the cytoskeletal proteins is observed in GFP control cells.

We have shown above that AdOx treatment has a significant effect on FUS condensate formation, but AdOx is also known as a drug that reduces DNA methylation. We therefore investigated the effect FUS condensates might have on the level of euchromatin formation, which is strongly increased by hypomethylation. We have recently shown that the dye SiR-DNA (or SiR-Hoechst) can be used to determine the level of eu/heterochromatin formation in live cells (Hockings et al., 2020; Novo et al., 2022). As SiR-DNA intercalates with DNA structures, its fluorescence lifetime is significantly affected by the level of chromatin condensation. Since euchromatin formation is associated with significant DNA decondensation, an increase in the fluorescence lifetime of SiR-DNA can be associated with an increase in euchromatin formation (Hockings et al., 2020; Novo et al., 2022). We observe that both FUS mutant cells contain more euchromatin (i.e., more decondensed DNA), with P525L-FUS cells (3.44±0.03 ns) experiencing a greater extent of DNA decondensation than R495X-FUS cells (3.43±0.04 ns) and in comparison to the GFP control (3.40±0.03 ns) and WT-FUS cells (3.41±0.03 ns) (**Figure 3C&D**). To further control our results, we applied two different drugs known to hinder DNA compaction, i.e., AdOx (due its effect on DNA hypomethylation (Schwerk and Schulze-Osthoff, 2005)) and Trichostatin A (TSA, which induces histone diacetylase (HDAC) inhibition (Moreira et al., 2003; Vigushin et al., 2001)) to control cells, which also raised the fluorescence lifetime of SiR-DNA to 3.44±0.04 and 3.43±0.03 ns, respectively. It is interesting to note that the effect aberrant FUS condensates has on euchromatin formation is as severe as the one induced by drugs, such as AdOx and TSA. Taken together, cytoskeletal defects as well as changes in euchromatin state are most prevalent in mutant variants of FUS, especially P525L-FUS.

**Figure 3:**
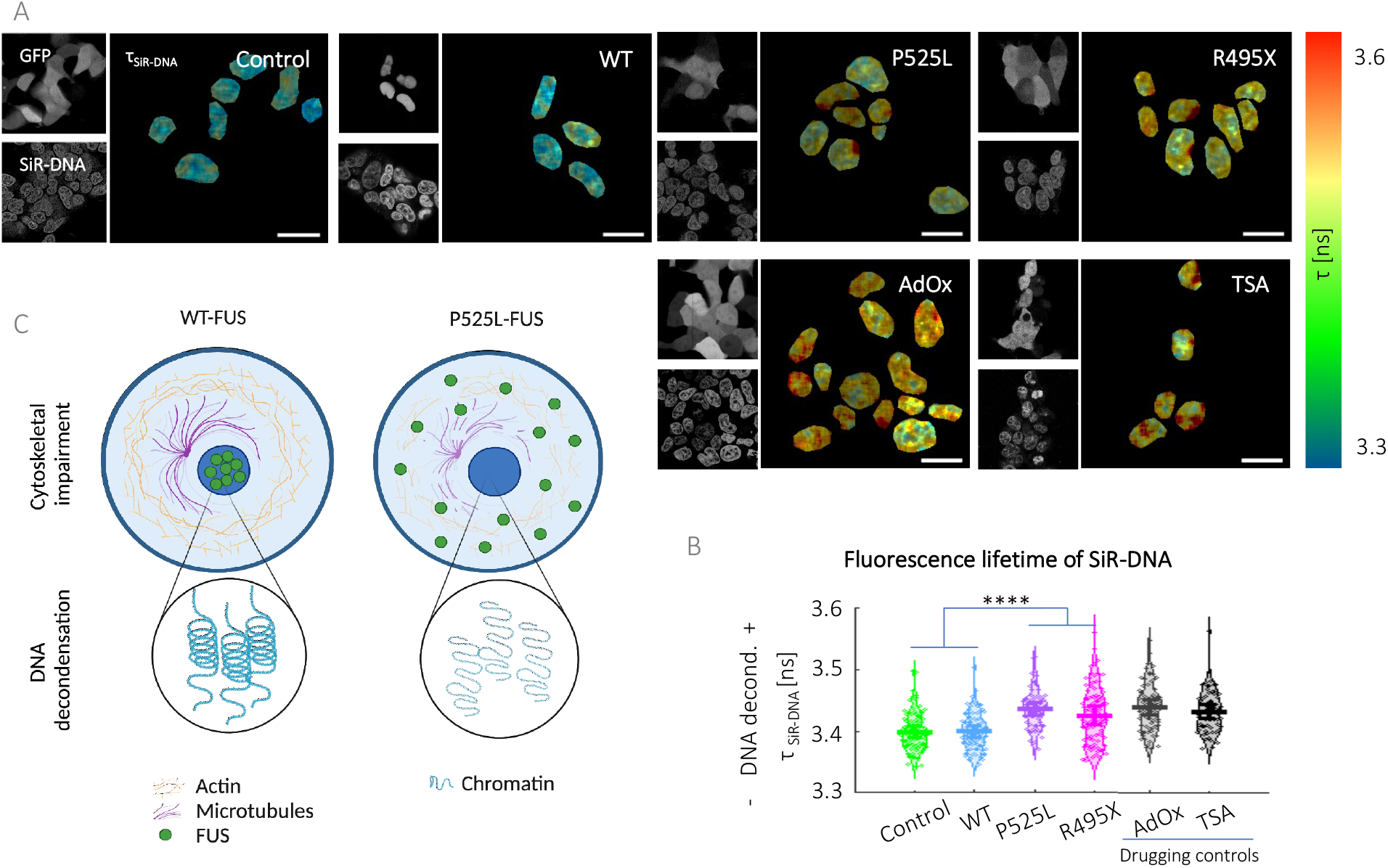
ALS-related FUS mutants lead to an increase in euchromatin formation. (A) Fluorescence lifetime of SiR-DNA is used as a measure of DNA decondensation, which is related to heterochromatin formation. Mutant FUS cells exhibit lower level of heterochromatin formation (i.e., higher τ_SiR-DNA_). GFP control cells treated with AdOx and TSA which cause DNA hypomethylation and HDAC inhibition, were included. (B) Fluorescence lifetime maps. Scale bars, 10 μm. (C) Cartoon illustration of impaired mechanoproperties of P525L-FUS cells due to weakened cytoskeleton and DNA damage in comparison to WT-FUS cells. Created on Biorender.com. HEK293T measurements were based on 3 biological repeats. One-way ANOVA test (with Holm-Sidak’s multiple comparison), where * for p<0.05, ** p<0.01, *** p<0.001, **** p<0.0001.

### Aberrant FUS aggregation impacts normal organelle function

The above studies have shown that mutant FUS has a significant effect on the cytoskeleton, which, in particular for tubulin. We thus investigated whether this effect on the microtubules might interfere with organelles, such as lysosomes, mitochondria, and ER, as the microtubules are important for the positioning, formation, fission, and fusion of the latter (Mattenberger et al., 2003; Pu et al., 2016). We applied super-resolution 2-colour structured illumination microscopy (SIM) with a resolution of 100 nm (i.e., below the diffraction limit) (Young et al., 2016) in COS7 cells, which have a flat morphology ideal for SIM, as the technique is sensitive to out-of-focus glare associated with thicker samples (Ma et al., 2021). We show that clustering of lysosomes and mitochondria at the perinuclear area of cells is greater in P525L-, compared to WT-FUS and control cells (**Figure S6**). We have recently found that lysosomes are responsible for the maintenance of the tubular ER network in the periphery of cells (Lu et al., 2022, 2020). Thus, clustering of lysosomes in the perinuclear area may significantly impair the tubular ER network in the cell periphery. Hence, to investigate if a collapse in the tubular ER structure also follows upon intracellular FUS accumulation, we co-expressed an ER marker, mEmerald-sec61β (Nixon-Abell et al., 2016) in mCherry (mChe) versions of our control and FUS cells. Synthetic, hypomethylation-mimicking variants which have increased numbers of arginine (i.e., 9R and 16R) were additionally included as further controls (Qamar et al., 2018). We observe that there is a significant impact on the tubular ER network as indicated by a decrease in the ER tubular to sheet ratio compared to the control upon FUS accumulation (**Figure S7A&B**). Furthermore, we also observe a decline in mitochondrial eccentricity (i.e., more rounded structures) in the presence of FUS accumulation, and particularly in P525L-FUS when compared to the control (**Figure S7D&E**), which may indicate defects in mitochondrial fission and increased mitochondrial stress.

Since lysosomes also play an important role not only the formation of the ER tubular network but also in protein homeostasis, we analysed their structure and function in the different FUS cells in more detail. To visualise the nanoscale structures of lysosomes in live cells, we again use SIM in mChe(-FUS) COS7 cells (**Figure 4A**, green in Composite). We additionally labelled lysosomes with LysoTracker Deep Red (**Figure 4A**, magenta in Composite), and segmented individual lysosomes from SIM images using a custom-written script (**Figure 4A**, where each lysosome is represented by a different colour). As a positive control, we treated mChe only control cells with chloroquine diphosphate (CQ). As an autophagy inhibitor, CQ promotes the formation and accumulation of autophagosomes, as it disrupts autophagy by preventing lysosome-autophagosome fusion(Mauthe et al., 2018). We observe that R495X (0.28±0.10 μm^2^ and 71±12 counted lysosomes) and P525L-FUS (0.29±0.12 μm^2^ and 62±9 counted lysosomes) have larger and more numerous lysosomes in comparison to WT- (0.12±0.04 μm^2^ & 46±5 counted lysosomes), 9R- (0.15±0.05 μm^2^ & 40±6 counted lysosomes) and 16R-FUS (0.14±0.03 μm^2^ & 46±7 counted lysosomes), which in turn are larger than in the control (0.09±0.03 μm^2^ and 23±3 counted lysosomes) (**Figure 4B&C**). As expected, the effect is most prominent in the CQ-added control sample (0.61±0.17 μm^2^ and 78±14 counted lysosomes).

**Figure 4:**
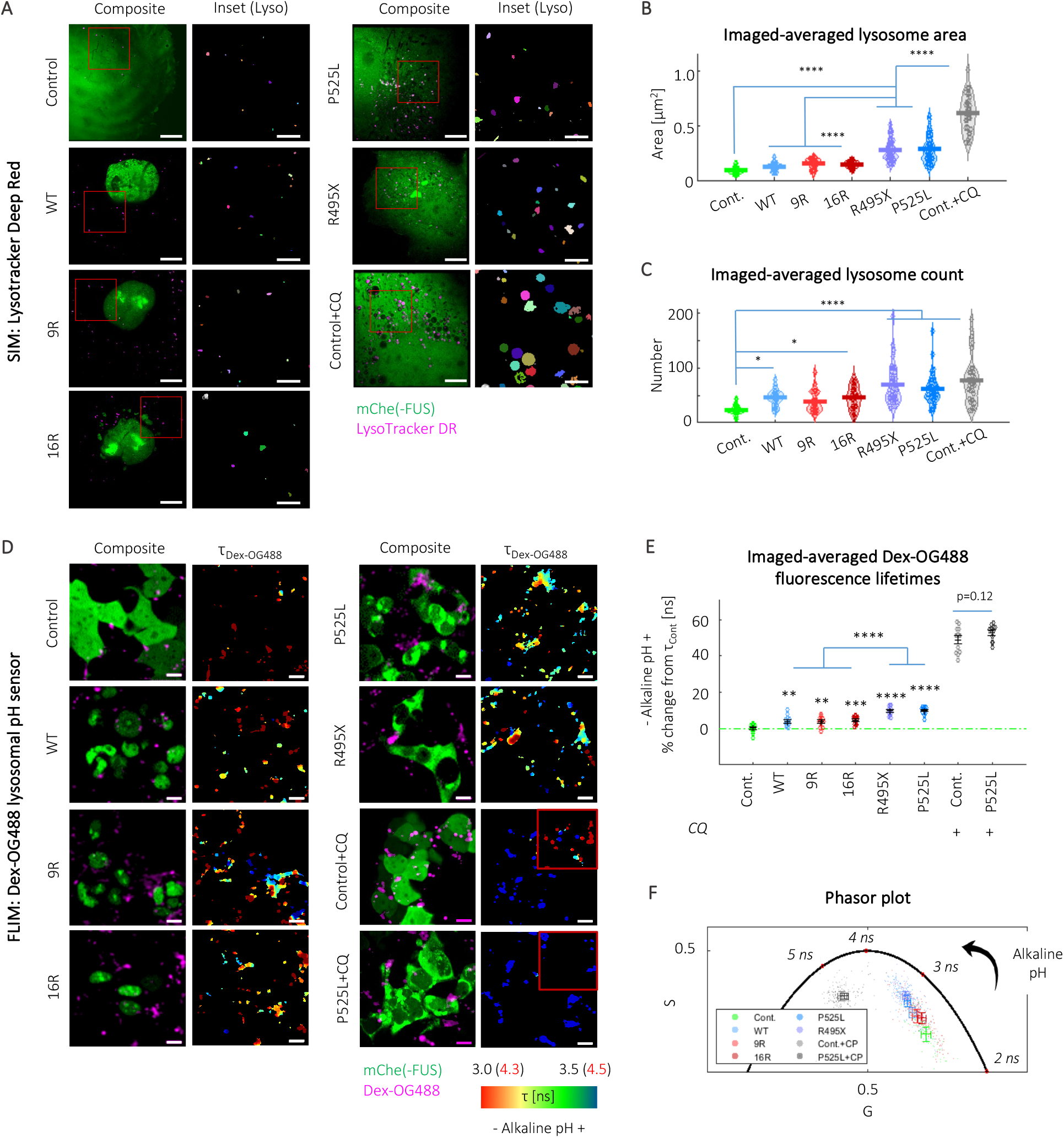
ALS-associated FUS mutants increase lysosomal size and biogenesis, and impact lysosomal pH. (A—C) SIM measurements to visualise lysosomal structure using LysoTracker Deep Red staining. (A) Composite SIM images showing mChe(-FUS) in green and LysoTracker Deep Red in magenta (first column, Composite). Inset corresponds to region within the red box in the Composite image, where individually segmented lysosomes are represented by a different colour. Scale bar, 10 μm. (B) Averaged lysosome area and (C) averaged lysosome count, which show that NLS-mutants, P525L and R495X have more, and Control+CQ has significantly more dilated lysosomes, in comparison to the control and nuclear-localised FUS variants. (D—F) FLIM measurements on Dex-OG488 lysosomal pH sensor. (D) Composite fluorescence intensity images showing mChe(-FUS) in green and Dex-OG488 in magenta (first column, Composite). The latter is used as a non-invasive lysosomal pH sensor with fluorescence lifetime-based readouts shown as falsely coloured images (second column, τ_Dex-OG488_). Scale bar, 10 μm. (E) Fit-free phasor plot showing fluorescence lifetime measurements for cell samples, where anti-clockwise direction indicates higher fluorescence lifetime and more alkaline pH. (F) Quantification of fluorescence lifetimes. Analysis is based on 12 images over three biological repeats. Analysis is based on 30—57 cells over three biological repeats. One-way ANOVA (with Holm-Sidak’s multiple comparison test), where n.s. is not significant, ** is p<0.005, *** is p<0.001 and **** is p<0.0001.

To test, whether these lysosomes are still able to maintain their low physiological pH, the latter of which is essential to maintain protein homeostasis, we employ Dextran 10,000 MW tagged to Oregon Green 488 (Dex-OG488), which has been used as a pH sensor in a fluorescence lifetime-based assay (Burdikova et al., 2015). Dex-OG488 accumulates in acidic compartments including lysosomes within cells (**Figure 4D**, Composite), hence can be used as an *in situ*, non-invasive lysosomal pH sensor. We measured Dex-OG488 fluorescence lifetime (τ_Dex-OG488_) using FLIM and analysed resulting data using fit-free phasor plot analysis. We see modest increases in τ_Dex-OG488_ for the nuclear-localised FUS variants, i.e., 3.92±0.99%, 3.67±0.95% (9R) and 4.31±0.72% (16R), in comparison to control cells, which indicates a rise in lysosomal pH. The differences become more significant for the NLS-mutants, i.e., 9.61±0.84% (R495X) and 9.66±0.67% (P525L), as well as CQ-treated control and P525L-FUS (48.4±2.4% and 52.4±1.44%, respectively) (**Figure 4E&F**). This is unsurprising as CQ is known to accumulate within lysosomes as a deprotonated weak base, and thereby increases lysosomal pH (Chen et al., 2011). In summary, we see that the dilated lysosomes in ALS-associated P525L- and R495X-FUS have lost their functionality due to greater lysosomal de-acidification.

### The presence of aberrant FUS aggregation promotes the nuclear translocation of TFEB but not autophagy

We have observed that greater lysosomal de-acidification leads to increased lysosomal biogenesis, both of which are enhanced in mutant P525L- and R495X-FUS. De-acidification of lysosomes may trigger autophagy via the nuclear translocation of TFEB, which is a transcriptional modulator for the autophagy-lysosomal pathway (ALP). The latter has been identified as a therapeutic target for diseases involving lysosomal dysfunction (Tan et al., 2021; Willett et al., 2017). Under physiological conditions, TFEB is phosphorylated and remains in the cytoplasm. Nuclear translocation and activation of the TFEB pathway are usually triggered under conditions of starvation and lysosomal stress, leading to autophagosome formation, and increased lysosomal biogenesis that upregulates autophagy (Settembre et al., 2011). Highly expressed in the central nervous system, the dysfunction of TFEB has been implicated in the pathogenesis of numerous neurodegenerative diseases including ALS/FTD (Chen et al., 2015). We thus addressed the role of the TFEB when it translocates from the cytoplasm to the nucleus. Upon co-transfection with GFP-TFEB into mChe(-FUS) HEK293T cells, we observe that there is greater nuclear translocation of TFEB when FUS is co-expressed, which in turn is amplified in the NLS-FUS mutants, P525L and R495X (**Figure 5A&B**). We quantify this as the ratio of nuclear to cytoplasmic fluorescence of GFP-TFEB from widefield images (**Figure 5C**, I_Nuc_/I_Cyto_), and validate that the observed effect is predominantly due to FUS expression, as all FUS-expressing cells have I_Nuc_/I_Cyto_ (e.g., 1.3±0.23 to 1.73±0.27 for WT- and P525L-FUS, respectively) that are significantly increased compared to the control cells (0.79±0.17). Control cells were also treated with CQ, which is a known stressor for lysosomal dysfunction, hence result in the induction of nuclear translocation of TFEB (Roczniak-Ferguson et al., 2012a) As expected, Control+CQ gives the greatest I_Nuc_/I_Cyto_ of 2.35±0.31 with 98% of cells showing nuclear translocation (i.e., I_Nuc_/I_Cyto_ >1); the latter is similar to both R495X and P525L at 86% and 90%, respectively.

**Figure 5:**
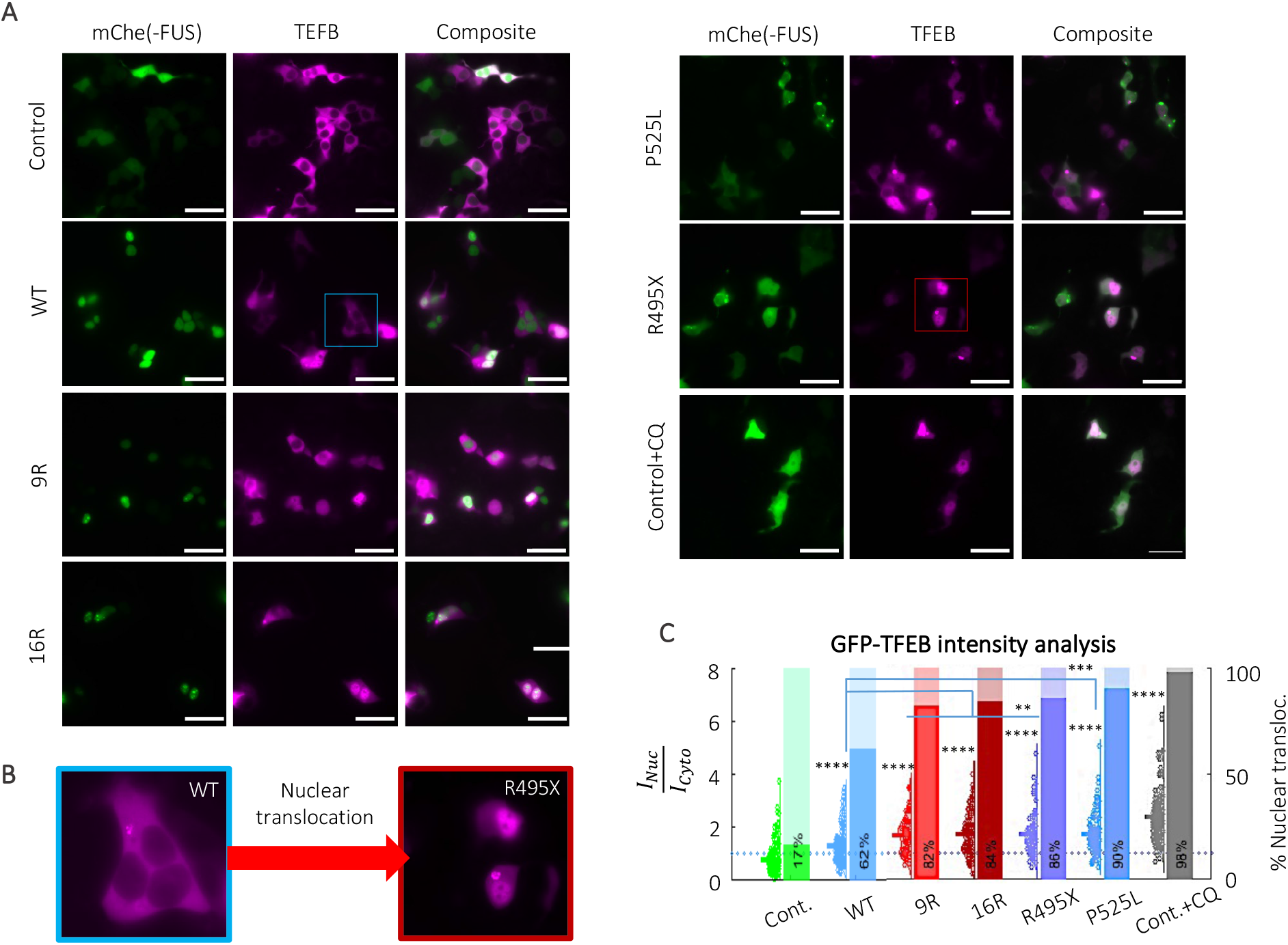
Nuclear translocation of TFEB is promoted by FUS accumulation, and in particular for ALS-associated mutants. (A) Widefield fluorescence intensity images showing mChe(-FUS) in green, GFP-TFEB in magenta and their composite image for all samples tested. Scale bar, 10 μm. (B) Images showing cytoplasmic TFEB expression in WT-FUS, and nuclear translocation in ALS-associated R495X-FUS. (C) Quantification of nuclear to cytoplasmic GFP-TFEB intensities (violin plot, primary y-axis), and their corresponding value of nuclear translocation (bar chart, secondary y-axis), based on images in (A). Mean and S.E.M. values of I_Nuc_/I_Cyto_: 0.79±0.17 (Control), 1.31±0.23 (WT), 1.68±0.23 (9R), 1.72±0.26 (16R), 1.70±0.23 (R495X), 1.73±0.27 (P525L) and 2.35± 0.31(Control+CQ). Analysis is based on 62—200 cells over three biological repeats. One-way ANOVA (with Holm-Sidak’s multiple comparison test), where n.s. is not significant, ** is p<0.005, *** is p<0.001 and **** is p<0.0001.

We have shown that the presence of FUS accumulation in cells could act as a stressor leading to a loss in acidification of lysosomes, the latter of which trigger the cytoplasmic to nuclear translocation of TFEB, as indicated by the increase in lysosomal biogenesis observed. To address if autophagy is initiated beyond lysosomal biogenesis, we performed Western Blot analysis for autophagy markers, i.e., p62 and LC3 (**Figure S8A&B**). With FUS accumulation, we observe a slight upregulation of p62, but not of LC3-II, both of which indicate that autophagy is inhibited at this early stage. The most significant difference occurs upon addition of CQ to the control, which drives up normalised LC3-II and p62 by 2- and 5-fold, respectively (**Figure S8B**). Furthermore, a FLIM-based aggregation assay was performed on both GFP-labelled control and P525L-FUS cells, treated with CQ, and rapamycin (i.e., an autophagy inducer). Of the latter, a relatively low concentration of 20 nM introduced in serum-starved media was used, as higher concentrations led to cell death. Fluorescence lifetime of the GFP tag remains the same for P525L-FUS, indicating that there are no changes in the aggregation state of FUS within the cells (**Figure S8C&D**). We thus show that autophagy, and therefore the clearance and degradation of FUS (and other proteins) are inhibited, despite activation of the TFEB-lysosomal pathway. Moreover, at this stage, rapamycin is either toxic to cells at higher doses or not efficacious in reducing FUS aggregation at lower concentrations.

## Discussion

The experimental data described here reveal that intracellular FUS accumulation leads to i) cytoskeletal and organelle dysfunction and ii) perturbations in cellular homeostasis. To study the latter, we not only analysed different cellular FUS and stress-related models with unprecedented detail but also developed an approach with which we could determine the level of viscosity in live cells.

From a micro-rheological standpoint, the liquid condensed state of FUS is a viscoelastic Maxwell fluid, possessing both viscous liquid and elastic solid behaviour (Jawerth et al., 2020). As part of this study, we have developed an approach, based on correlative SPT and FLIM to measure intracellular FUS viscosity. This provides a higher throughput method than FRAP, commonly used in this field, as several condensates can be imaged simultaneously. Moreover, this approach allows us to probe condensates which are smaller than those suitable for more conventional FRAP analysis. It provides a robust route to quantifying this intracellular viscosity, for which there is no existing tool that is widely applicable in the field. We find that NLS-mutants of FUS, P525L- and R495X-FUS, contain more viscous aggregates in comparison to WT-FUS cells. Of the two FUS mutant variants studied, P525L-FUS has the greater aggregation propensity, leading to higher accumulation of drug-induced intranuclear aggregates and SGs. We further show that these mutant variants are also associated with a loss in cytoskeletal mechanoproperties in the cytoplasm, as seen by significant impairment of their actin and tubulin networks. As part of our study, we used SiR-dyes which permit this effect to be captured in live cells, as current studies typically rely on immunofluorescence staining of fixed samples. We further corroborated this using non-fluorescence-based, physical FD-AFM measurements, which show that P525L has half the apparent Young’s modulus value of WT-FUS. Similar observations in neuronal models have linked axonal cytoskeletal integrity to the proteinopathy of FUS (Castellanos-Montiel et al., 2020; Giampetruzzi et al., 2019; Theunissen et al., 2021) and other FTD/ALS-implicated proteins, e.g., TAR DNA-binding protein (TDP43) (Baskaran et al., 2018; Briese et al., 2020). This includes a related study in *Xenopus laevis* retinal ganglion cells that demonstrated P525L-FUS reduces actin density specifically in growth cones, resulting in localised softening of the axon(van Tartwijk et al., 2022). Furthermore, several ALS-linked genes that directly impair cytoskeletal dynamics (Castellanos-Montiel et al., 2020; Garone et al., 2020). We further observe that NLS-FUS mutant cells undergo DNA decondensation. At physiological conditions, nuclear-localised FUS regulates DNA repair through its interactions with HDAC1, which becomes dramatically reduced in the case of mutant FUS (Naumann et al., 2018; Wang et al., 2013; Yang et al., 2014) and can in turn increase euchromatin formation (Dinant et al., 2008; Lukas et al., 2011; Miller et al., 2010). Since epigenetic and chromatin regulatory factors are also phase separating in the nucleus (Gibson et al., 2019; Larson et al., 2017; Sanulli et al., 2019; Strom et al., 2017), it is conceivable that mutant, and thus more viscous FUS in the nucleus may also affect epigenetic and chromatin regulatory factors and thereby further enhance euchromatin formation.

We, among others, have recently shown that the dysfunction of common biochemical pathways which attenuate organelle function (e.g., lysosomes, ER, mitochondria), are prevalent to multiple neurodegenerative disease(Fowler et al., 2019; Koh et al., 2019; Lu et al., 2020). In combination, they lead to a loss in cellular integrity, and eventually culminate in neuronal dysfunction. To address organelle distribution and function in various cellular FUS and stress-induced models, we applied several different imaging techniques and assays. With ALS-associated mutant P525L-FUS, we observe that increased lysosomal biogenesis leads to lysosomal clustering, as well as mitochondria clustering, at the perinuclear region of cells. As a result, we see a decline in tubular ER due to the loss in lysosomal distribution throughout the cell cytoplasm. Tubular ER is predominantly found in axons, where it modulates synaptic calcium for neurotransmitter release (de Juan-Sanz et al., 2017), and supplies membranes and proteins for synaptic function(Gómez-Suaga et al., 2019). Hence any disruption to its function may have significant consequences for neurons. Additionally, the tubular ER also regulates mitochondrial fission (Merkwirth and Langer, 2008), and thus the mitochondria clustering seen in FUS-expressing cells may influence the latter. P525L-FUS has the largest effect on mitochondria shape, as indicated by the significant loss of eccentricity, i.e., the mitochondria become more circular from a tubular structure, which may stem from the effect of FUS on mitochondrial fission.

Many of the above results point towards a dysregulation of the ALP which is supported by previous studies showing aberrant FUS aggregation disrupts the latter (Baskoylu et al., 2022; Soo et al., 2015). We observe cytoplasmic to nuclear translocation of TFEB in the presence of FUS accumulation, which we relate to the higher intralysosomal pH measured in FUS cells. In association, we observe increased lysosomal biogenesis in the absence of changes to autophagy levels. These results support the key role TFEB plays in autophagy regulation, and hence the adverse effects its dysfunction yields. Despite using non-neuronal HEK293T and COS7 cells, we believe the underlying mechanisms observed should hold true for a more physiologically relevant model. In support of this, it was recently shown that there is impaired autophagy and neuronal dysfunction in a P525L-FUS knock in *C. elegans* model (Baskoylu et al., 2022). Moreover, our findings align with the fact that autophagy dysfunction occurs early on in disease pathology, which acts as an accelerator for neurodegeneration (Cortes and la Spada, 2019). Autophagy becomes even more vital in the case of aged, i.e., non-/slow-dividing neurones. It has been shown that autophagy-deficient mice are more likely to accumulate aggregation-prone proteins, increasing the risk of neurodegeneration (Hara et al., 2006; Komatsu et al., 2006). Proper function of TFEB and lysosomes have been associated with ageing and longevity (Chang et al., 2017; Lapierre et al., 2015). A recent study has shown that promotion of lysosomal function through the overexpression of Pep4 (i.e., Cathepsin D homologue) in a yeast cell model leads to extended lifespans (Carmona-Gutiérrez et al., 2011). Interestingly, our and the above findings contrast with the interactions of TDP43 and TFEB, where it has been shown that the nuclear translocation of TFEB occurs with a deficiency in the first and leads to increased lysosome/autophagosome biogenesis (Xia et al., 2016). However, the loss in TDP43 also leads to downstream impairment of lysosome-autophagosome fusion, leading to an accumulation of autophagic vesicles, and thus both TDP43 and FUS models display autophagy dysfunction. Further studies will be required to pinpoint the exact mechanisms of TFEB-lysosomal dysfunction as well as its impact on organelles, in the presence of aberrant FUS aggregation. Interestingly, many of the defects we have observed as part of our study pinpoint towards an induction of early senescence, such as loss of cellular mechanoproperties, increased euchromatin formation, and loss of protein homeostasis. Aberrant phase transition of FUS had been hypothesised to drive cellular ageing by others in the field (Alberti and Hyman, 2016). It thus remains to be determined whether neuronal cells indeed undergo early senescence as part of a cellular stress induced by aberrant FUS and whether the latter precedes the formation of FUS condensates.

There is no current therapeutic strategy for ALS/FTD and many other neurodegenerative diseases. Autophagy inducers, e.g., rapamycin, have been proposed as treatments to alleviate aberrant FUS aggregation (Zhang et al., 2011). However, in our cell models, we did not observe any alleviation in FUS aggregation upon rapamycin treatment. We have previously proposed that perinuclear lysosomal clustering leading to the collapse of the ER network could be a hallmark mechanism across neurodegenerative disease (Lu et al., 2020). Moreover, this has a further impact on cellular integrity and function. For FUS, this is additionally seen in the depolymerisation of cytoskeletal proteins. Hence, potential therapeutic strategies may want to be targeted towards alleviating cytoskeletal defects, organelle dysfunction and perturbations in cellular homeostasis, rather than solely towards aberrant protein aggregation.

Our study paints a picture of the effect of aberrant FUS condensates which is highly complex, and impacts many key cellular structures/functions, such as the cytoskeleton, organelles, and cellular homeostasis, which in turn, cannot be alleviated by targeting only one of these defects. Interestingly, many of the above defects mimic signs of early senescence. Indeed, similar features resembling cellular senescence have also been described in other neurodegenerative (Martínez-Cué and Rueda, 2020; Sahu et al., 2022). It remains to be determined whether aberrant proteins first trigger a stress response that affects major cellular functions, such as related to cellular senescence, before or after the formation of aberrant condensates/aggregates. Understanding the latter, will have a major impact on the understanding common factors underlying neurodegenerative diseases and for the development of new therapeutic approaches.

## Materials and Methods

### Preparation of recombinant FUS condensates

For both SPT and FLIM measurements, equal amounts of GFP and SNAP-tagged recombinant WT-FUS protein (yielding a total protein concentration of 6 μM) were gently mixed into an aqueous solution of 20 % PEG-35 (Merck KGaA, Darmstadt, Germany) and 500 mM KCl (Merck KGaA) in a protein lo-bind tube (Eppendorf, Hamburg, Germany). Both recombinant proteins were a gift from the Alberti group (Max Planck Institute of Molecular Cell Biology and Genetics, Dresden, Germany). For the formation of fibrils, 20% PEG-35 was omitted from the mixture, and the solution was incubated for 20 mins at room temperature before imaging. Recombinant WT-FUS proteins were a gift from the Alberti group (Max Planck Institute of Molecular Cell Biology and Genetics, Germany). For SPT, the aqueous solution also included a 10^−5^ dilution of 40 nm fluorescence nanoparticles (FluoSpheres carboxylate-modified microspheres, red-orange fluorescence, ThermoFisher Scientific, Waltham, MA, USA), and the solution was sonicated for 15 minutes before each use. 7 μL of the condensate mixtures was deposited in a silicon well (Press-to-Seal, ThermoFisher Scientific) attached on 1.5 thickness coverslips (Superior Marienfeld, Lauda-Konigshofen, Germany) for ageing and imaging.

### Cell culture

COS7 and HEK293T cells (American Type Cell Culture, Manassas, VA, USA) were cultured in T25/T75 cell flasks with media, were incubated at 37 °C and 5% CO_2_. Culture media comprised of 90 v/v% Dulbecco’s Modified Eagle’s Media (DMEM, ThermoFisher Scientific), 10 v/v% foetal bovine serum (FBS, ThermoFisher Scientific), and 2 mM each of glutamax (ThermoFisher Scientific) and 2% penicillin-streptomycin (ThermoFisher Scientific). Cells were passaged when 80–90% confluency was reached (i.e., twice a week). Cells were plated into 8 well plates (μSlide 8 Well, IBIDI GmBH, Gräfelfing, Germany) to achieve 70-80% confluency on the day of imaging. For hypomethylated intranuclear aggregates, 20 μM adenosine periodate, oxidase (AdOx, Merck KGaA, Darmstadt, Germany) were added to cell media 24 hours before imaging. 2.5 μM sodium arsenite (Merck KGaA) was added to cell media 90 minutes before imaging for stress granule formation.

### FUS expression plasmids & cell transfection

HEK293T stable cell lines expressing GFP only control as well as GFP-WT, P525L and R495X-FUS were synthesised using lentivirus-based constructs. FUS plasmids were a gift from Dr. S. Qamar (Cambridge Institute for Dementia Research (CIMR))(Qamar et al., 2018).

Cells were plated overnight in antibiotics-free media to achieve 40—70% confluency on the day of transfection. For 8 well transfections, 200 ng of DNA plasmid and 0.6 μL Lipofectamine 2000 reagent (ThermoFisher Scientific) were mixed into 60 μL OptiMEM (ThermoFisher Scientific) and incubated at room temperature for 20 minutes. The DNA-lipid mixture was then added into media of the desired well and incubated for 4—6 hours at 37°C. Media change into antibiotics-added media was performed after, and the cells were further incubated for 20 hours before imaging.

### Fluorescence lifetime imaging microscopy (FLIM)

Samples were imaged on a home-built confocal fluorescence microscope equipped with a time-correlated single photon counting (TCSPC) module. A pulsed, supercontinuum laser (Fianium Whitelase, NKT Photonics, Copenhagen, Denmark) provided excitation a repetition rate of 40 MHz. This was passed into a commercial microscope frame (IX83, Olympus, Tokyo, Japan) through a 60x oil objective (PlanApo 60XOSC2, 1.4 NA, Olympus). For GFP-labelled *in vitro* system and HEK293T cells, and OG488-Dex in HEK293T cells, the excitation and emission beams are filtered through GFP-appropriate bandpass filters centred at 474 and 542 (FF01-474/27-25, FF01-542/27, Semrock Inc., NY, USA). Laser scanning was performed using a galvanometric mirror system (Quadscanner, Aberrior, Gottingen, Germany). Emission photons were collected on a photon multiplier tube (PMT, PMC150, B&H GmBH, Berlin, Germany) and relayed to a time-corelated single photon counting card (SPC830, B&H GmBH). Images were acquired at 256×256 pixels for 120 s (i.e., 10 cycles of 12 s). Photon counts were kept below 1% of laser emission photon (i.e., SYNC) rates to prevent photon pile-up. TCSPC images were analysed using an in-house, MATLAB-based (MathWorks, Natnick, MA, USA) phasor plot analysis script (https://github.com/LAG-MNG-CambridgeUniversity/TCSPCPhasor), from which fluorescence lifetime maps and phasor plots were generated. FLIM results presented are based on 3 biological repeats for HEK293T cells and 3 individual protein preparations for recombinant FUS condensates.

### Single particle tracking (SPT) in recombinant FUS condensates

The widefield microscope uses an IX83 (Olympus) frame, and excitation light is provided by a 4-wavelength LED source powered by a DC4100 driver (Thorlabs, Newton, NJ, USA). GFP and red-orange fluorescence FluoSpheres were imaged using microscope filter cube sets for GFP and m-Cherry, through a 60x oil immersion objective lens (PlanApo 60XOSC2, 1.4 NA, Olympus) and a Xyla sCMOS camera (Andor, Belfast, UK). For SPT trajectories, 1000 frames were captured at a frame rate of ∼50 frames per second.

SPT analysis was used to calculate the intra-condensate viscosity. Nanoparticles within condensates were located, and their coordinates were tracked over the time frames. Neighbouring coordinates were linked together based on their time points in the image sequence, yielding a 2-dimensional trajectory of the tracked nanoparticle. Adopting a passive micro-rheology approach, it is assumed that random motion of the nanoparticle is due to thermal fluctuations.(Crocker and Grier, 1996) Mean square displacement (MSD) profiles of individual nanoparticles were extracted. Stokes-Einstein equation (Equation 1) was used to calculate viscosity (μ).

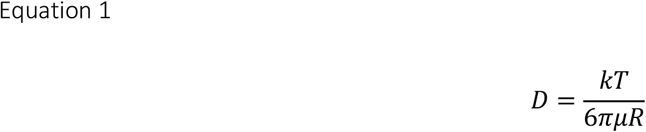

where D is diffusion coefficient, k is the Boltzmann’s constant, T is temperature (i.e., 298 K), R is nanoparticle radius (i.e., 20 nm).

### Immunofluorescence staining of G3BP in fixed HEK293T

Cell media was removed and replaced with 4% paraformaldehyde (PFA, Merck KGaA) diluted in phosphate buffer solution (PBS, ThermoFisher Scientific). The sample was fixed for 15 minutes, before permeabilising the cells with 0.1 w/v% Triton X-100 (ThermoFisher Scientific) and blocking them with 5 w/v% bovine serum albumin (BSA, Merck KGaA), both diluted in PBS for 1 hour. Between antibody incubation, three washes of 50 μM Triton X-100 (Invitrogen, ThermoFisher Scientific) in PBS (henceforth referred to as PBST) were performed. Primary anti-G3BP antibody (ab56574, abcam) was diluted at 1:200 in PBST and the cells were incubated at 4 °C overnight. Alexa Fluor 647 labelled secondary antibody (goat anti-rabbit IgG, Alexa Fluor 647, ThermoFisher Scientific) was used at a dilution of 1:400 and cells were incubated for 1 hour with the antibody at room temperature. The sample was kept wrapped in aluminium foil to prevent any bleaching especially after the addition of the secondary antibody and stored at 4°C.

### Staining and confocal imaging of the cytoskeleton in HEK293T

SiR-fluorogenic dyes for microtubules and f-actin (Spirochrome, Thurgau, Switzerland) were used for staining live cells at 1 μM alongside 10 μM verapamil (Spirochrome), followed by an incubation period of 30 minutes at 37°C. For all stains described in this subsection, a single wash, followed by addition of fresh DMEM. Imaging was performed using the same confocal microscope setup as that of the TCSPC-FLIM. For SiR-dyes, supercontinuum excitation was passed through a bandpass excitation and emission filters centred at 632 and 700 nm respectively (FF02-632/22-25, Semrock Inc; ET700/75m Chroma, Burlington, VT, USA). Emission photons were detected using a single photon counting module avalanche photodiode (SPCM-AQRH, Excelitas Technologies, Mississauga, Canada), and images were acquired at 1024×1024 pixels. Manual masks were drawn for individual cells in each image. A MATLAB script then calculated the mean intensity values of SiR-tubulin or actin within each cell.

### Force-displacement (FD-) atomic force microscopy

Live cells were imaged in phenol red free DMEM (ThermoFisher Scientific) in 35 mm AFM imaging cell dishes. FD-AFM measurements were performed on a BioScope Resolve (Bruker GmbH, Karlsruhe, Germany), under PeakForce QMN mode using a silicon nitride tip of 70 nm in diameter (PFQNM-LC-A-CAL, Bruker GmbH). Images were collected at a scan rate of 1 Hz and resolution of 256×256 pixels. For each individual FD curve, baseline correction was performed, followed by fitting to a linear Hertzian model with a Poisson ratio of 0.5, on NanoScope Analysis 9.4 (Bruker GmbH). Output values were imported to MATLAB in a .csv format. An automated script that selectively accepts apparent Young’s modulus values calculated based on linear fitting with a coefficient of determination (R^2^) above 0.8, was used to process the data. Results were performed over 3 biological repeats.

### DNA decondensation assay using SiR-DNA

SiR-DNA (Spirochrome) was performed using the same protocols as detailed for SiR-Tubulin and SiR-Actin. For GFP controls, cells were treated with 20 μM AdOx (Merck KGaA) and 200 ng/mL Trichostatin A (TSA, ThermoFisher Scientific) for 24 hours and 60 minutes respectively, before imaging. TCSPC imaging and analyses were performed as previously described.

### Calculating diffusion coefficient of FUS macroassemblies in live cells

Live cells were imaged after 90 mins of NaAsO_2_ and 24 hours of AdOx treatment on the same widefield setup as for SPT imaging. Recordings of hypomethylated aggregate movements were capture across 1000 frames, and their diffusion coefficients were found using analysis detailed under SPT.

### Overexpression, widefield imaging and analysis of TFEB

GFP-TFEB plasmids(Roczniak-Ferguson et al., 2012b) were transfected into cells using Lipofectamine 2000 (as detailed in the previous subsection). For imaging, the widefield microscope used is based on a IX83 (Olympus) frame, and excitation light is provided by a 4-wavelength LED source powered by a DC4100 driver (Thorlabs, Newton, NJ, USA). GFP-TFEB and mCherry-FUS were imaged using microscope filter cube sets for GFP and m-Cherry, through a 60x oil immersion objective lens (PlanApo 60XOSC2, 1.4 NA, Olympus) and a Xyla sCMOS camera (Andor, Belfast, UK). Manual masks for nuclear and cytoplasmic regions were drawn for the cells imaged. Corresponding intensity values for these areas were quantified using an in-house, automated MATLAB (MathWorks, Natnick, MA, USA) script, which also calculated nuclear to cytoplasmic intensity ratio values.

### Western blot

HEK293T cells grown in 6 well plates were lysed by pipetting up and down in radioimmunoprecipitation assay (RIPA) buffer with added protease inhibitors (Pierce, ThermoFisher Scientific), and kept overnight at -80°C. Upon thawing on ice, cell lysates were sonicated using an ultrasonic bath (UH-300, UltraWave, Cardiff, UK) for three 1-minute runs over 6 minutes. Protein concentration was measured using a bicinchoninic acid (BCA) assay (Pierce BCA Protein Assay Kit, ThermoFisher Scientific), and 20 mg was boiled alongside sample loading buffer (Invitrogen NuPAGE LDS 4X, ThermoFisher Scientific) for 5 minutes at 95°C. All following reagents used were purchased from Invitrogen NuPAGE (ThermoFisher Scientific), unless otherwise stated. Samples were loaded into a pre-cast gel wells (10%, Bis-Tris, 15 well, 1 mm), with pre-stained protein ladder (PageRuler, 10 to 180 kDa, ThermoFisher Scientific), for electrophoresis in MES SDS running buffer (1x diluted in PBS from 20X). Protein transfer onto a polyvinylidene difluoride (PVDF) transfer membrane (0.45 μm, ThermoFisher Scientific) was performed in transfer buffer (1x diluted in PBS from 20X). Electrophoresis and gel transfer were performed using a XCell SureLock Mini Cell Electrophoresis System (Invitrogen ThermoFisher Scientific). The PVDF membrane was blocked in 5 w/v% milk (skim milk powder for microbiology, Millpore, Merck KGaA) in PBS for 1 hour on an orbital shaker (SC5, Stuart Equipment, ThermoFisher Scientific). The resulting membrane was then cut to allow for primary antibody staining of β-tubulin (ab15568, abcam, Cambridge, UK), f-actin (ab130935, abcam), LC3 (ab192890, abcam) and p62 (ab109012, abcam); and the housekeeping gene, PCNA (ab18197, abcam) overnight at 4°C on a tube roller (SRT6, Stuart Equipment). Secondary antibody staining using either sheep α-mouse or donkey α-rabbit (NA931V, NA934V, GE HealthCare, Chicago, IL, USA) was performed for 1 hour on a tube roller at room temperature. Three 5-minute washes in 0.1 v/v% Tween-20 (Merck KGaA) in PBS was included after both primary and secondary antibody staining.

The blot was developed using SuperSignal West Pico PLUS Chemiluminescent Substrate (ThermoFisher Scientific) between two sheets of acrylic and imaged on a G-box (Chemi XX6, Syngene, Bengaluru, India). Quantification of protein bands was performed using a custom MATLAB script which normalises the intensity sum of each over that of PCNA.

### Organelle staining/labelling in live cells

COS7 cells were used for all SIM experiments. mEmerald-Sec61β-C1 plasmids(Nixon-Abell et al., 2016) were transfected into cells using Lipofectamine 2000 (as detailed in the previous subsection). On the day of imaging, mitochondria and lysosomes were stained using MitoTracker Deep Red and LysoTracker Deep Red (ThermoFisher Scientific), at final concentrations of 200 nM and 50 nM for 30 minutes at 37°C. Dextran, Oregon Green 488 (10,000 MW, Anionic, ThermoFisher Scientific) was added to FUS-expressing cells at a final concentration of 1.25 μg/mL. They were left to incubate overnight at 37°C. Cells were washed once in fresh media before imaging.

### Structured illumination microscopy (SIM) and analysis

The SIM used is a home-built system that uses a spatial light modulator (SLM) for generating SIM grating patterns, as described in Young *et al*.(Young et al., 2016). Imaging was performed using a water immersion objective lens (UPLSAPO 60 XW, 60x/1.2 NA, Olympus). For GFP(-FUS), mCherry(-FUS) and MitoTracker Orange, and Mito/LysoTracker Deep Red excitation, 488 nm (iBeam SMART, Toptica, Munich, Germany), 561 nm (OBIS LS, Coherent, Santa Clara, CA, USA) and 647 nm (MLD, Cobolt AB, Stockholm, Sweden) diode lasers were used respectively. Emission light were passed through respective bandpass filters (FF01-525/30, FF01-676/29, Semrock) onto a sCMOS camera (ORCA Flash 4.0, Hamamatsu, Shizuoka, Japan) at exposure times between 10— 100 ms. SIM reconstruction was performed using fairSIM, an open-source FIJI plugin (Müller et al., 2016).

### Segmentation and morphological analysis of organelles

Batch and automated analysis were performed using a custom MATLAB script, which reads images of both channels of 2-colour SIM images as input. On the image corresponding to the organelle of interest, an intensity threshold is set to separate background from desired features. A clustering algorithm performs segmentation based on defined size criteria to ensure that individual organelles (i.e., single mitochondria or lysosome) are segmented. Ambiguous segmented objects are eliminated from further analysis. For each segmented organelle, a size (e.g., area and major length axis) and morphological (e.g., eccentricity) quantification is performed.

### Statistical analyses and plotting

All statistical analyses were performed on Prism 6 (GraphPad, San Diego, CA, USA), where one-way ANOVA test with Holm-Sidak’s multiple comparison were applied. Results are given as n.s. for not significant, * for p<0.05, ** p<0.01, *** p<0.001, **** p<0.0001. Violin plots were produced by adapting open-source MATLAB code from Anne Urai (github.com/anne-urai).

## Acknowledgements

CWC is funded by the Cambridge Trust and Wolfson College for her PhD. GSKS acknowledges funding from the Wellcome Trust (065807/Z/01/Z) (203249/Z/16/Z), the UK Medical Research Council (MRC) (MR/K02292X/1), Alzheimer Research UK (ARUK) (ARUK-PG013-14), Michael J Fox Foundation (16238) and Infinitus China Ltd. PHStG-H acknowledges funding from the Canadian Institutes of Health Research (406915 Foundation Grant and Canadian Consortium on Neurodegeneration in Aging Grant), Wellcome Trust Collaborative Award (203249/Z/16/Z), US Alzheimer Society Zenith Grant (ZEN-18-529769), Alzheimer Society of Ontario Chair in Alzheimer’s Disease Research and National Institute of Aging (U01AG072572; R01AG070864).

## Contributions

Study conceptualisation and design: GSKS; Cell culture and plasmids: CWC, ADS & AM; Data collection: CWC, AJZ & IM; Data analysis: CWC & AJZ; Feedback: PHStG-H, CFK & TJPK; Draft manuscript preparation: CWC & GSKS; Manuscript editing: all authors. All authors have given approval of the final manuscript.

## Declaration of interests

The authors declare no competing interests.

## Supporting information

Methodology and supplementary figures (PDF)

## Data availability

Raw data is available through Cambridge University Repository Apollo (DOI: 10.17863/CAM.83295). Analysis scripts are available upon request.

## Notes

https://www.doi.org/10.17863/CAM.83295

### Competing Interest Statement

The authors have declared no competing interest.

### Summary of Updates

Reference update

