## Supporting Information for "Intracellular FUS protein accumulation leads to cytoskeletal, organelle and cellular homeostasis perturbations"

### Supplementary Figures

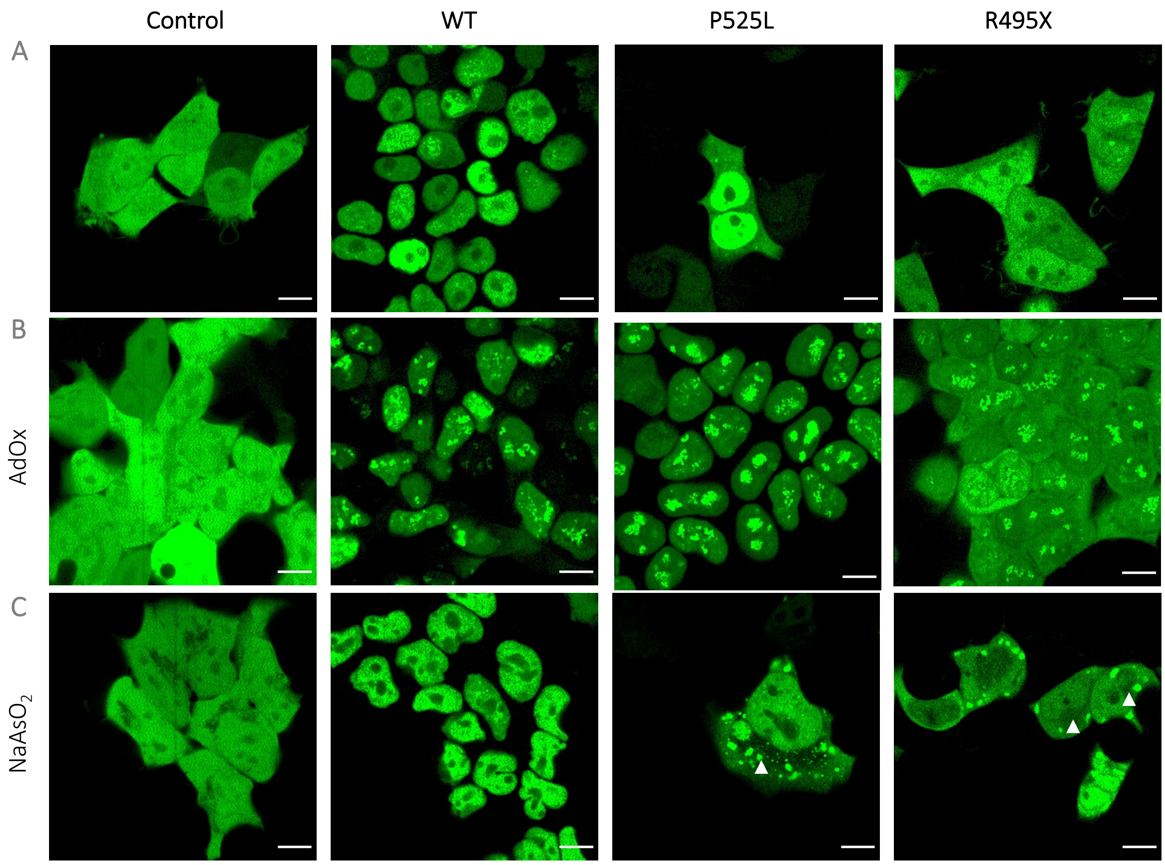

**Figure S1: Mutant FUS causes aberrant cytoplasmic localisation of FUS, whereas stress induced by AdOx and $NaAsO_2$ causes increased nuclear condensates in all FUS positive cells and cytoplasmic stress granules in mutant FUS cells, respectively.** (A) In contrast to the GFP-only control, WT-FUS cells show nuclear FUS localisation, in contrast to NLS-mutant, P525L- and R495X-FUS cells. To induce the formation of FUS macro-assemblies, we treated them with stressors: (B) AdOx or (C) $NaAsO_2$, which lead to the formation of hypomethylated aggregates and stress granules (the latter only in mutant FUS as highlighted by white arrows). Scale bar, 10 µm.

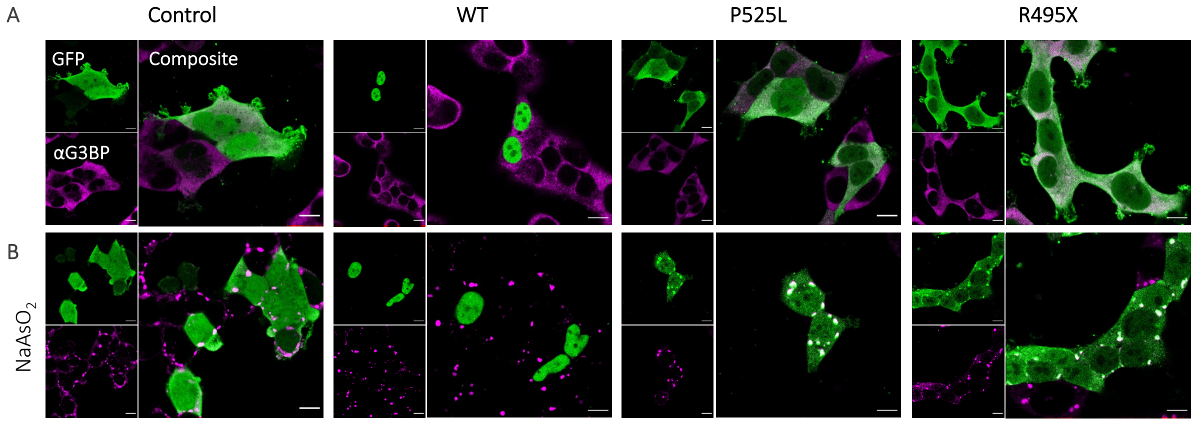

**Figure S2: Aberrant FUS aggregation within SGs only occurs in the case of mutant FUS.** GFP(-FUS) and G3BP (a SG marker)-positive cells are shown in green and magenta respectively. (A) G3BP is dispersed throughout the cytoplasm. (B) Addition of stressor, $NaAsO_2$ triggers the formation of SGs; however, aberrant FUS aggregation (i.e., colocalisation of FUS and G3BP) is only seen in mutant P525L and R495X-FUS cells. Scale bar, 10 µm.

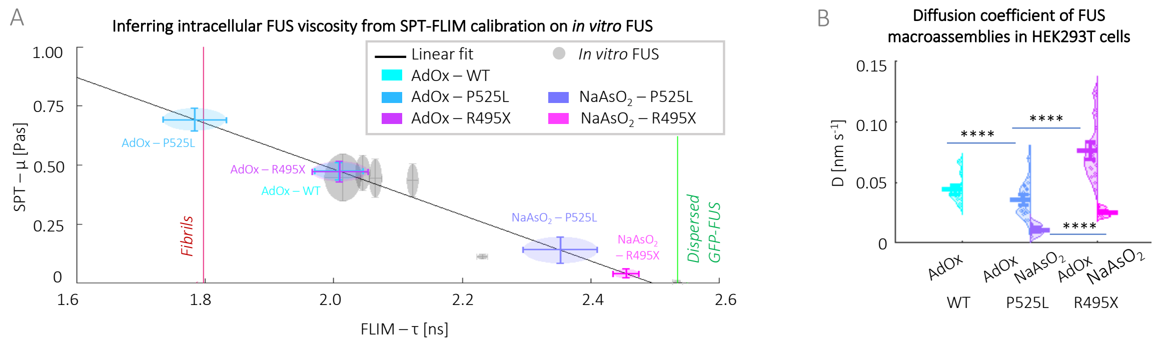

**Figure S3: Macroassemblies of P525L-FUS have greater viscosity and lower mobility, implying greater aggregation and more severe disease progression.** (A) Fluorescence lifetime to viscosity calibration based on *in vitro*-formed condensates (grey circles) show a linear relationship (black line), onto which measured fluorescence lifetimes of whole cells (without treatment) and different macroassemblies formed upon treatment, of different FUS variants are mapped. AdOx-treated cells have viscosity values ~0.5 Pas, which is several folds larger than that of SGs at ~0.05 Pas. (B) Diffusion coefficient (D) of FUS macroassemblies. P525L-FUS cells have the least mobile nuclear aggregates and SGs, in comparison to WT and R495X-FUS cells.

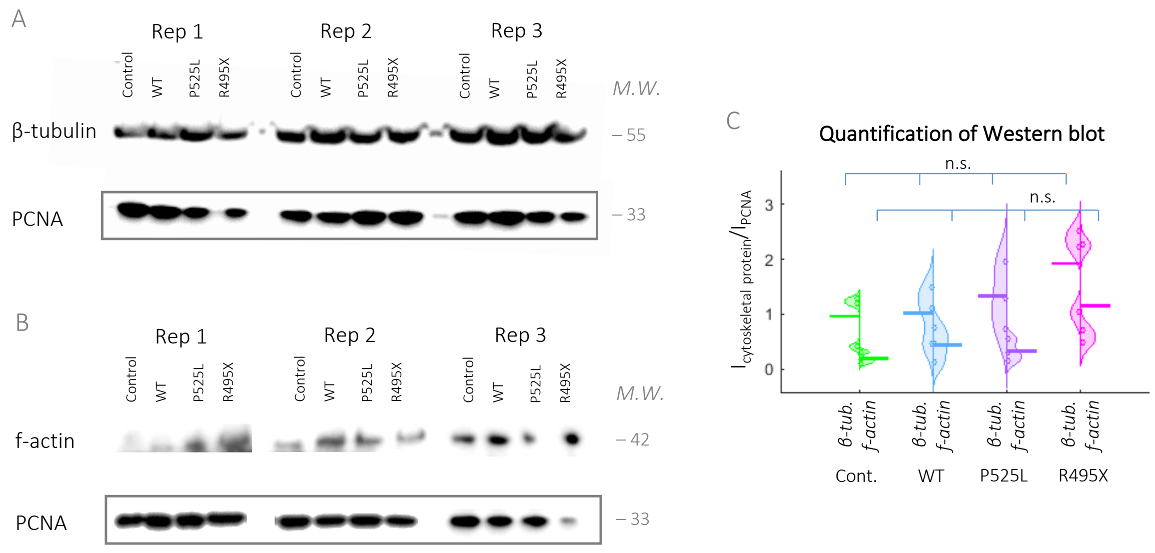

**Figure S4: Western blot shows that total concentration of f-actin and β-tubulin in cells do not differ significantly.** (A—B) Western blot of 3 different biological repeats (Rep) of (A) f-actin, (B) β-tubulin and PCNA (i.e., housekeeping) expressions from lysed cells. Note that f-actin and β-tubulin analysed on individual blots. (C) Quantification of (A&B) showing that expression of both β-tubulin and f-actin is not significantly different.

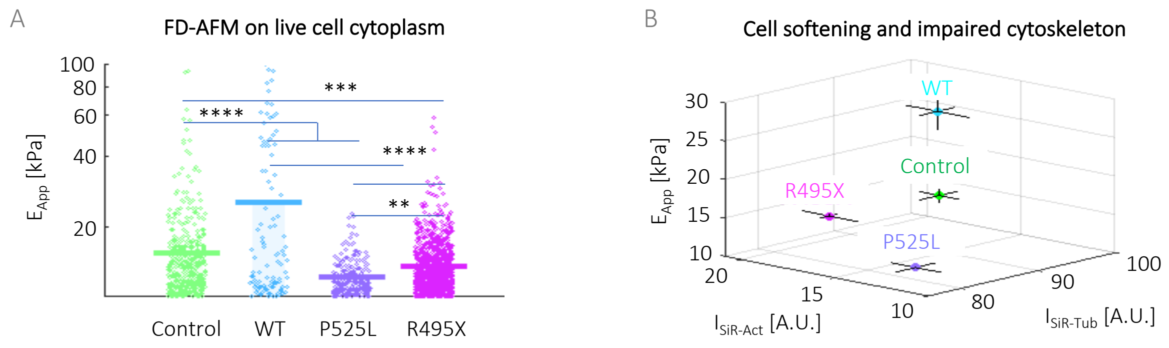

**Figure S5: ALS-related FUS mutants impair cytoskeletal mechanoproperties.** (A) Quantification of apparent Young's moduli ($E_{App}$) by FD-AFM performed on live cells, which show that P525L and R495X-FUS cells have a softer cytoplasm in comparison to WT-FUS cells. (B) Cell softening can be correlated to loss in cytoskeletal proteins, the latter is based on SiR-Actin and Tubulin staining shown in **Figure 2**.

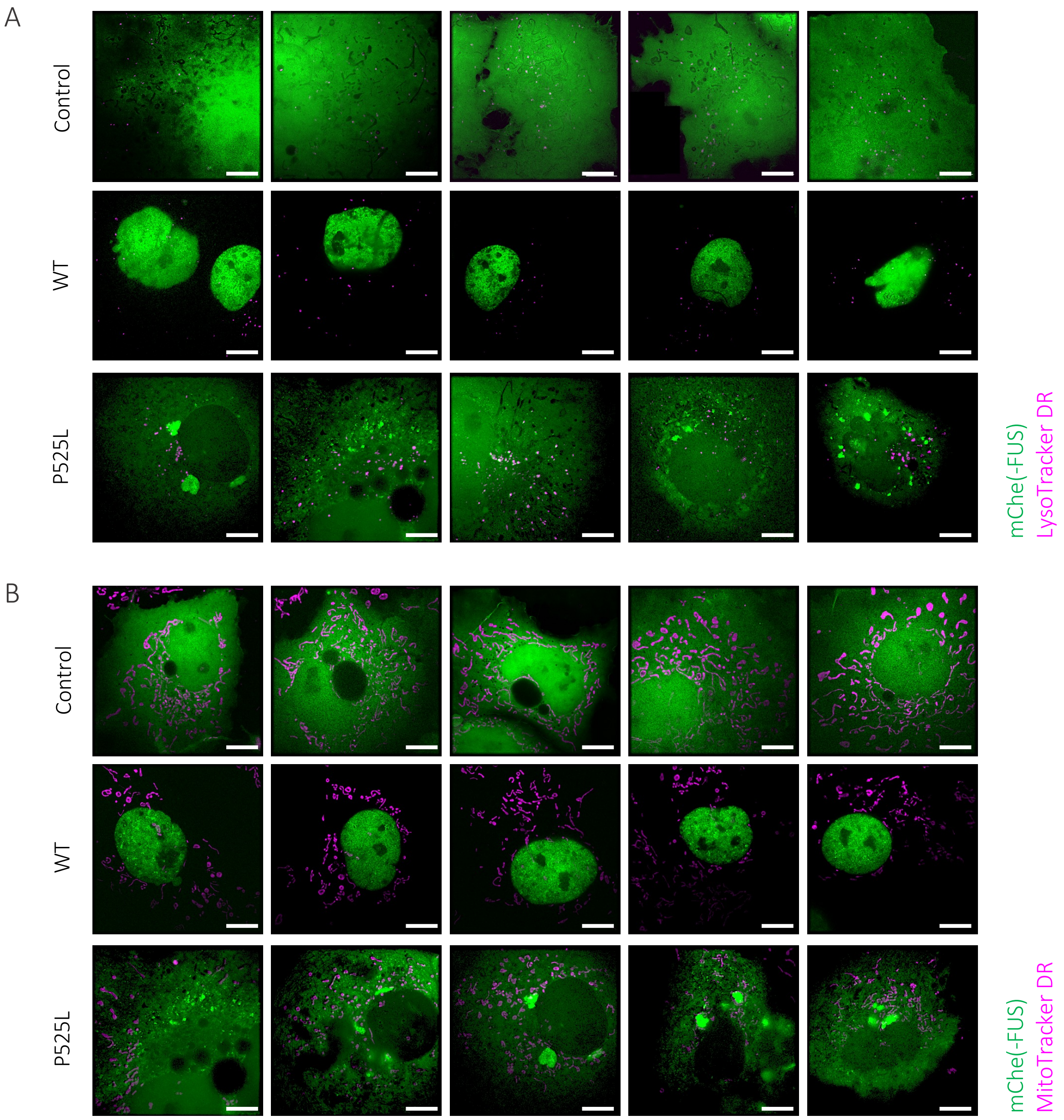

253 **Figure S6: Greater perinuclear accumulation of lysosomes and mitochondria for P525L-FUS.** Composite SIM images showing mChe(-FUS) with
254 either (A) lysosomes labelled with LysoTracker Deep Red or (B) mitochondria with MitoTracker Deep Red. Scale bar, 10 µm.

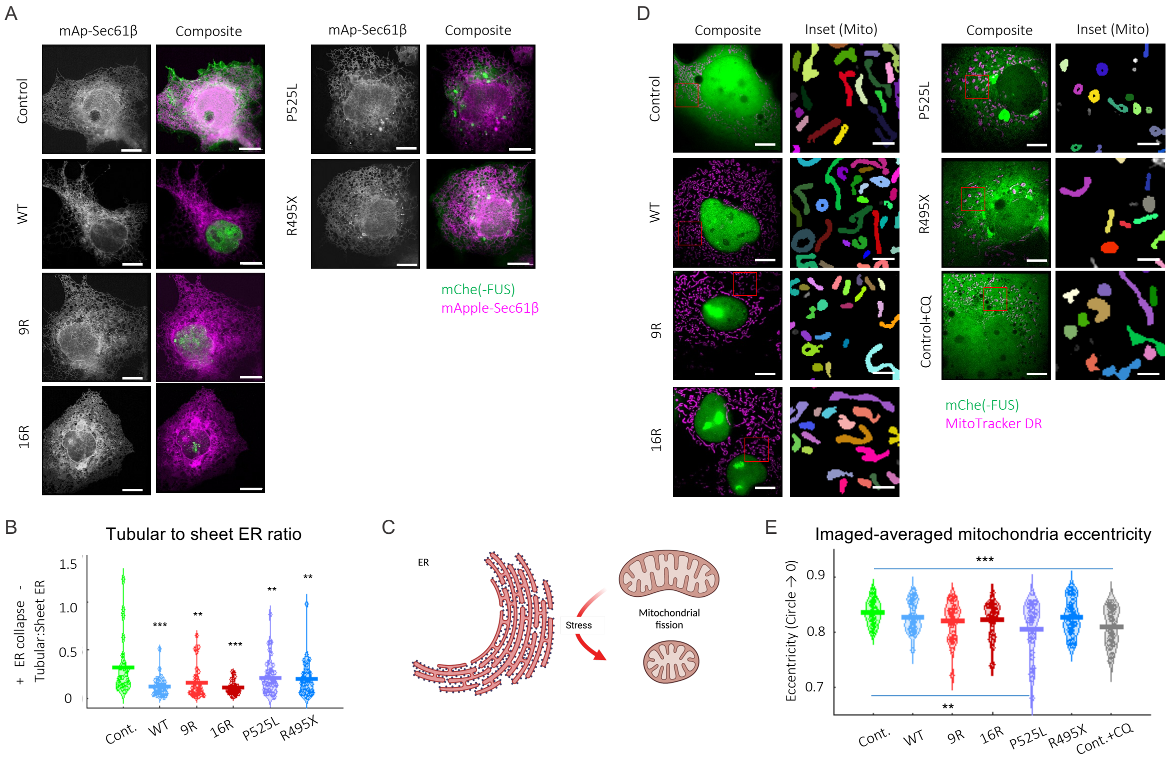

**Figure S7: Reduction in tubular ER network leads to greater mitochondrial fission with FUS-expression.** (A) SIM images showing ER-marker, mApple-Sec61β (first column) and composite images where mChe(-FUS) is in green and the ER is in magenta. (B) Quantification of tubular to sheet ER ratio using ERNet. (C) Cartoon schematic showing the loss in tubular ER leads to greater mitochondrial fission (i.e., more rounded mitochondria). (D) SIM images showing mChe(-FUS) in green and MitoTracker Deep Red-labelled mitochondria in magenta. Inset shows individually segmented mitochondria, corresponding to region within the red box in the Composite image. (B) Quantification of mitochondrial eccentricity, showing that ALS-associated mutant P525L and CQ-treated control cells have rounder mitochondrial morphology, indicating greater mitochondrial fission. Scale bars, 10 µm. Results based on 26—39 cells taken over 3 biological repeats. One-way ANOVA (Holm-Sidak's multiple comparisons test), where n.s. is not significant, ** is p<0.005, *** is p<0.001 and **** is p<0.0001.

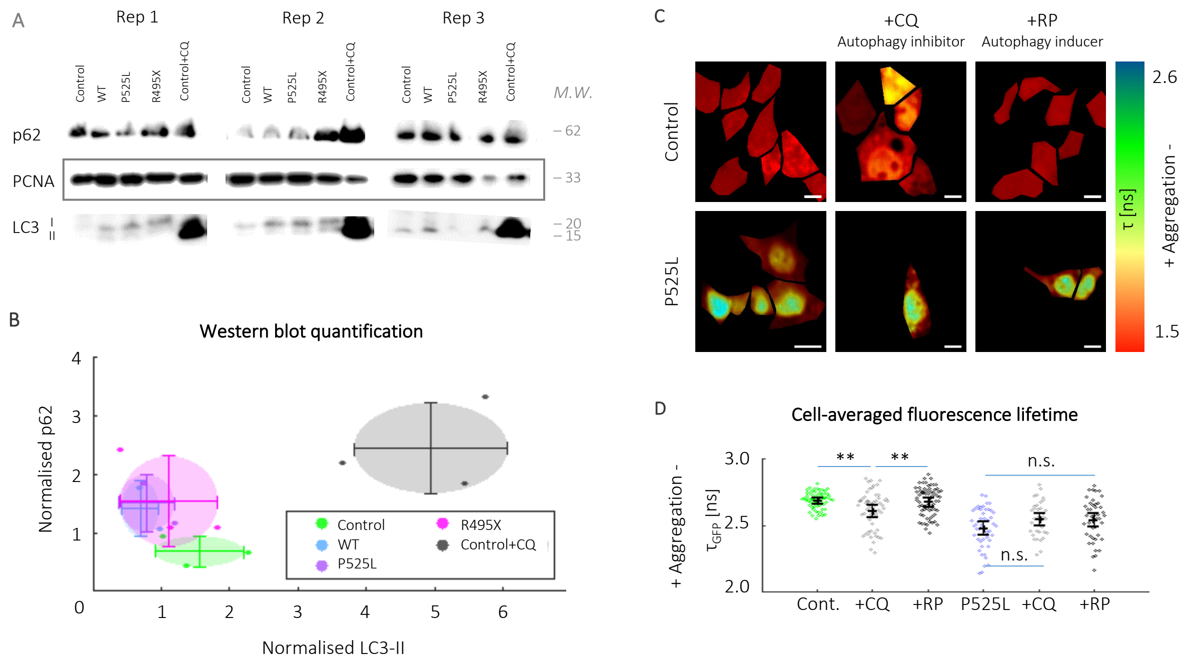

**Figure S8: Increased autophagy is not seen in FUS-expressing cells and P525L-FUS aggregation is not alleviated by low concentration rapamycin treatment.** (A) Representative Western blot showing p62, LC3 and the housekeeping gene PCNA. (B) Quantification of Western blot band intensity for LC3-II and p62, normalised to PCNA. Only Cont.+CQ (grey) is significantly different from the other cell samples. Analysis based on 3 biological repeats. (C) Fluorescence lifetime maps of GFP only (Control) and GFP-P525L cells treated with 2.5 µM chloroquine diphosphate (CQ, i.e., autophagy inhibitor) and 20 nM rapamycin (RP, i.e., autophagy promoter) overnight (the latter under serum-starved conditions). (D) Quantification of GFP fluorescence lifetime, where there is no difference before and after either drug treatments. Results based on over 30 cells taken from 3 biological repeats. One-way ANOVA (Holm-Sidak's multiple comparisons test), where n.s. is not significant, ** is $p<0.005$, *** is $p<0.001$ and **** is $p<0.0001$.
